# A corticothalamic circuit trades off speed for safety during decision-making under motivational conflict

**DOI:** 10.1101/2021.11.21.469477

**Authors:** Eun A Choi, Medina Husić, E. Zayra Millan, Philip Jean Richard dit Bressel, Gavan P. McNally

## Abstract

Decisions to act while pursuing goals in the presence of danger must be made quickly but safely. Premature decisions risk injury or death whereas postponing decisions risk goal loss. Here we show how mice resolve these competing demands. Using microstructural behavioral analyses, we identified the spatiotemporal dynamics of approach-avoidance decisions under motivational conflict. Then we used cognitive modelling to show that these dynamics reflect the speeded decision-making mechanisms used by humans and non-human primates, with mice trading off decision speed for safety of choice when danger loomed. Using calcium imaging and functional circuit analyses, we show that this speed-safety trade off occurs because increases in paraventricular thalamus (PVT) activity increase decision caution, thereby increasing approach-avoid decision times in the presence of danger. Our findings demonstrate that a discrete brain circuit involving the PVT and its prefrontal cortical input dynamically adjusts decision caution during motivational conflict, trading off decision speed for decision safety when danger is close. They identify the corticothalamic pathway as central to cognitive control during decision-making under conflict.

---

Foraging animals balance the need to seek food and energy against the conflicting needs to avoid injury and predation. Resolving this conflict is fundamental to survival, but decisions about whether to approach or avoid a goal with conflicting value rarely have a stable solution. Instead, appropriate solutions vary across space and time (diurnal, seasonal) as well as internal states (hunger, thirst). Making these decisions requires matching sensory information about the world to knowledge about reward or danger to support adaptive behavior. These decisions must be made quickly and safely. They require balancing the competing demands of speed in decision-making with safety of choice. Premature decisions risk injury or death whereas failures to decide in a timely manner risk goal loss. Although much is known about the brain mechanisms of danger and reward (Mobbs et al., 2020), the mechanisms for this decision-making under motivational conflict, including those controlling the trade-off between decision speed and decision safety, are poorly understood.

Here we studied how mice make approach-avoidance decisions under motivational conflict. We studied the microstructure of behaviour under conflict and used cognitive modelling to identify the latent mechanisms of choice. We show that mice trade-off decision speed for decision safety when danger looms. We then examined how the spatiotemporal activity dynamics of two brain regions – the paraventricular thalamus (PVT) and ventral subiculum (vSub) which have been implicated in motivational conflict (Choi et al., 2019; Choi and McNally, 2017; Engelke et al., 2021; Gray and McNaughton, 2000; Ma et al., 2021; Zhu et al., 2018) - relate to these spatiotemporal variations in choice mechanics. We found that activity in PVT, but not vSub, closely tracks approach–avoidance decision-making during motivational conflict. We show that a discrete brain circuit involving the PVT and its prefrontal cortical input dynamically adjusts decision caution when making choices under motivational conflict. Our findings show that decision-making under motivational conflict in mice shares the same lawful features as speeded decision-making in human (Forstmann et al., 2016) and non-human primates (Shadlen and Shohamy, 2016) and we identify a novel role for PVT in cognitive control that that may extend across a range of motivated behaviours.

## Results

### Microstructure of behavior under conflict

We trained thirsty mice (n = 8) to run a linear track, obtaining a liquid sugar reward from a goalbox (**Figure 1a**). Then, conflict was introduced by electrifying the floor of the goalbox with current (0.05, 0.075, 0.1 mA) that increased across days. Under these conditions, mice learn spatial gradients of reward and punishment that they use to select unique approach or avoidance decisions across the track (Hull, 1938; Miller, 1944). As expected, conflict decreased running speeds, increased the time taken to obtain reward, and reduced reward consumption (**Figure S1, Table S1**).

**Figure 1.**
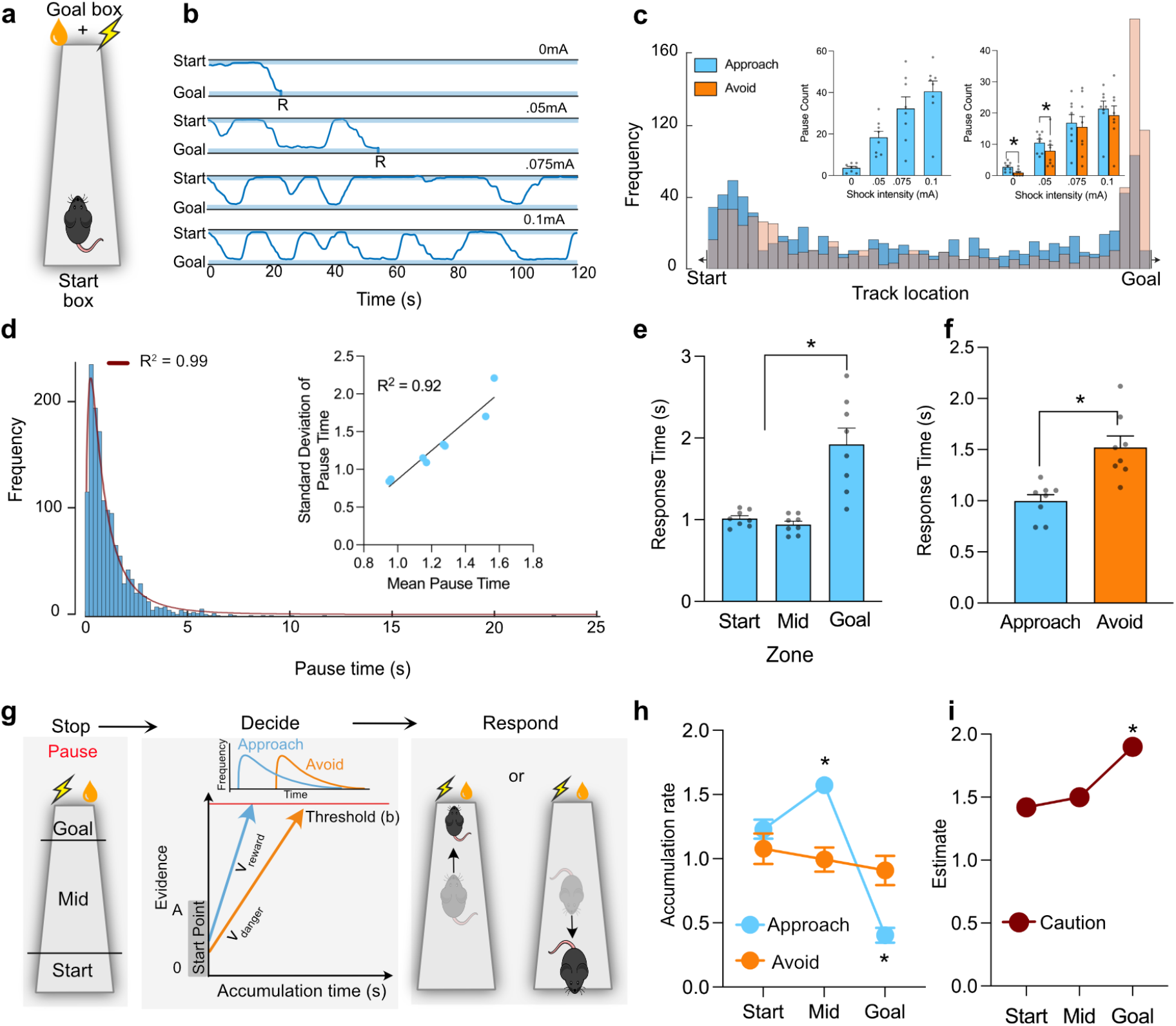
Spatiotemporal properties of decision-making under motivational conflict. **a**, Conflict paradigm. **b**, Example trajectories from no conflict (0 mA) and conflict tests. R indicates consumption of reward. **c**, Spatial distribution of pauses, inset shows mean and SEM (n = 8) pause counts, overall and by decision outcome. **d**, Frequency distribution of pause durations and log-normal fit. Inset shows correlation of mean pause times and standard deviations. **e**, Mean and SEM response times for the three track zones. **f**, Mean and SEM response times for approach versus avoid decisions. **g**, The Linear Ballistic Accumulator (LBA) model of choice. Evidence (V) for reward or danger accumulates separately to an evidence threshold (b) in a winner takes all process. **h**, Mean and SEM accumulation rates for approach and avoid decisions across the zones. **i**, Mean and SEM response caution across the zones. All * p < .05. See Table S1 for detailed results.

Visualization of individual trajectories (**Figure 1b, Figure S2**) showed that behavior under conflict rapidly developed bistable oscillations between the start and the end of the track. These oscillations were interrupted by pauses, ranging from milliseconds to seconds in duration. During these pauses, mice would rear, show lateral head scanning movements, and investigative sniffing (Thompson et al., 2018). These pauses were distributed across the track, but peaked at the start of the runway and just prior to the goal (**Figure 1c**), consistent with pauses reflecting active sampling of the environment to prompt individual approach or avoid decisions (Hull, 1938; Miller, 1944; Redish, 2016).

By identifying each pause, we could ask how each approach–avoid decision was resolved. There were notable spatial biases in decision outcomes. Approach decisions dominated across most of the track but were replaced by avoidance decisions closer to the goal (**Figure 1c**). In the absence of danger, there were few decisions, and these were resolved in favor of approach. In the presence of danger, the frequency of decision-making increased and the approach bias was lost (**Figure 1c inset;** shock intensity: *F* (3, 21) = 20.38, p = 0.0001; approach v avoid: 0mA *t* (7) = 3.230 p = 0.0144, 0.05mA *t* (7) = 2.582 p = 0.0363).

We measured response times (RTs) (i.e. time to choose approach or avoid after pause onset) because RTs are the most robust and widely used measure of decision-making efficiency (Laming, 1968). RTs were positively skewed, log-normal functions (R^2^ = 0.99, *F* (1, 122) = 1312, p < 0.0001) with linear coefficients of variation (R^2^ = 0.93, *F* (1, 6) = 80.59, p < 0.0001) (**Figure 1d**). These properties were preserved across choice outcome and levels of danger (**Figure S3**). We then used cluster analysis to ask if these properties depended on spatial location. This identified three zones of decision-making on the track (excluding start and goal box): start zone (the first 15cm after the start box), goal zone (last 5 cm of track), and mid zone (remainder of track). Approach-avoid decisions at the goal zone were most difficult, taking the longest time (main effect location: *F* (2, 14) = 24.86 p = 0.0014), start v goal: *t* (7) = 4.564 p = 0.026; start v mid: *t* (7) = 5.555 p = 0.0009) (**Figure 1e**). There was a trade-off between the speed of decision-making and the safety of choice. Avoid decisions leading to safety were more difficult (i.e. slower) than approach decisions (**Figure 1h**) (*t* (7) = 6.395 p = 0.0004).

These decision-making dynamics are shared with speeded choice in human (Forstmann et al., 2016) and non-human primates (Shadlen and Shohamy, 2016). They can be explained by sequential sampling models that decompose choice into its latent cognitive mechanisms (Brown and Heathcote, 2008; Forstmann et al., 2016; Ratcliff et al., 2016; Wagenmakers and Brown, 2007). Here, learned spatial gradients of reward and punishment are sampled from the environment and memory until an evidence threshold is reached and an approach or avoid choice made. The Linear Ballistic Accumulator (LBA) (Brown and Heathcote, 2008) is one such sequential sampling model that permits complete analytic solution for choices between alternatives (**Figure 1g**). We used Bayesian estimation via Hamiltonian Markov Chain Monte Carlo (HMC) to quantitatively fit the LBA to each animal’s approach-avoid decisions and derive key decision-making parameters: the rate of evidence accumulation for an approach decision (reward salience), the rate of evidence accumulation for avoid decision (danger salience), and the threshold of evidence required to reach a decision (response or decision caution).

The LBA fit the behavioral data well, explaining both RT distributions and choice outcomes (**Figure S4**). Parameter estimation showed that evidence accumulation and decision caution varied dynamically across the track. Reward salience increased across the track (Mid: *t* (7) = 8.134 p < 0.0001 vs avoid) but decreased significantly at the goal zone (*t* (7) = − 5.555 p = 0.0005 vs avoid) (**Figure 1h**), explaining the switch from approach to avoid decisions at the goal zone. In contrast, decision caution increased at the goal zone (*t* (7) = 4.245 p = 0.0038 versus mid zone) (**Figure 1i**), showing that more evidence was required to reach a decision as danger loomed and explaining the increased RTs near the goal.

### Paraventricular thalamus but not ventral subiculum tracks approach–avoidance decisions

Two brain regions, the ventral subiculum (vSub), located in the temporal lobe, and the paraventricular thalamus (PVT), located in the midline thalamus, have been implicated in motivational conflict (Choi et al., 2019; Choi and McNally, 2017; Engelke et al., 2021; Gray and McNaughton, 2000; Ma et al., 2021; Zhu et al., 2018), and so could contribute to these decision-making dynamics. Yet how activity in these brain regions relates to approach-avoidance decision-making is still poorly understood. To address this, we infused an adeno-associated viral vector (AAV) encoding the calcium (Ca^2+^) sensor gCaMP7 (Dana et al., 2019) into the vSub or PVT and implanted a fiber optic cannula above these regions. We then used fiber photometry to record vSub (n = 5) or PVT (n = 6) population Ca^2+^ signals of the mice during decision-making under conflict (**Figure 2a, Figure S5**).

**Figure 2.**
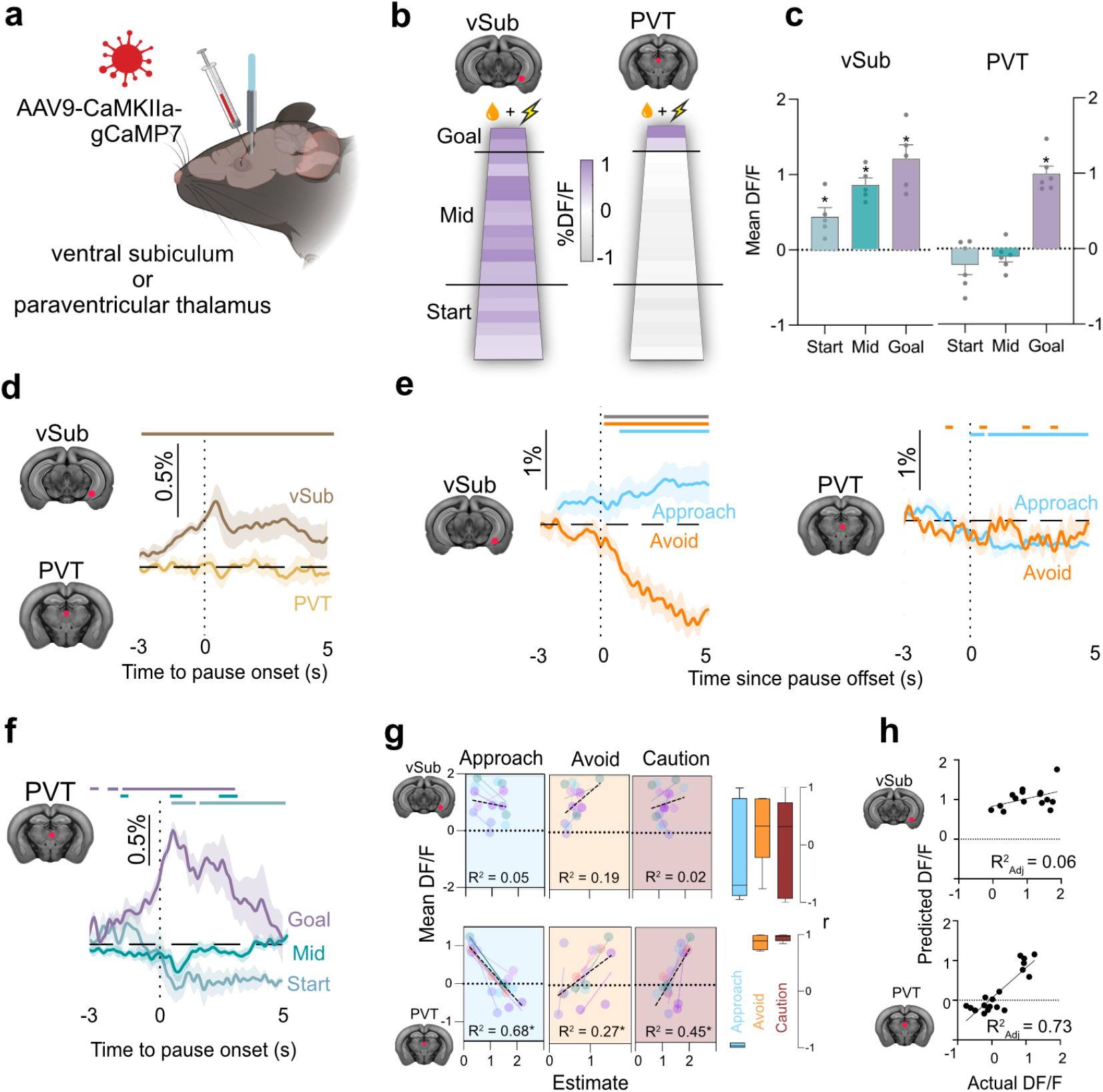
Paraventricular thalamus and ventral subiculum dynamics during conflict. **a**, AAV encoding gCaMP7F targeted to PVT or vSub. Fiber optic cannula implanted above injection site. **b**, Mean %DF/F across the track in 5 cm blocks during conflict (PVT n = 6, vSub n = 5). **c**, Mean and SEM %DF/F for Start, Mid, and Goal zone locations (excluding star and goal boxes). **d**, Mean and SEM %DF/F at pause onset (0 s) across the track. Colored bars indicate periods when 95% confidence interval (CI) does not include 0% DF/F. **e**, Mean and SEM %DF/F at pause offset (0 s) by decision outcome at Start zone. **f**, PVT mean and SEM %DF/F at pause onsets by location. Colored bars indicate periods when 95% CI does not include 0% DF/F, gray bar indicates significant difference between approach versus avoid via 95% CI. **g**, Relationship between LBA parameters for approach (reward salience), avoid (danger salience), response caution and mean %DF/F during pauses across track zones fitted by individual mouse (colored points and lines) and overall (black dotted line) with overall R^2^. *Right panel*, Box plots of individual mouse correlation coefficients. **h**, Linear model of PVT Ca^2+^ transients from LBA approach and avoid accumulation rates. * p < .05. See Table S2 for detailed results.

vSub Ca^2+^ transients were observed across the track (**Fig. 2b**), with activity increasing as mice approached the dangerous goal (**Fig. 2b,c**) (repeated measures ANOVA *F* (2, 12) = 7.51 p = 0.0077; versus 0% DF/F - start: *t* (4) = 3.620 p = 0.0224; mid *t* (4) = 9.147 p = 0.0008; goal *t* (4) = 6.413 p = 0.0030). Consistent with a role for vSub in behavioral inhibition during conflict (Gray and McNaughton, 2000), vSub activity increased prior to and during individual approach-avoid decisions across the track (**Figure 2d;** 95% Confidence Intervals). vSub activity also depended on choice outcome, remaining elevated if mice chose to approach the dangerous goal but not if they chose avoidance to the safety of the start box (**Figure 2e;** 95% Confidence Intervals).

In contrast, PVT Ca^2+^ transients showed a spatial bias, with significant increases in activity only at the goal zone (**Figure 2b,c**) (repeated measures ANOVA *F* (2, 15) = 40.62 p < 0.0001; versus 0% DF/F - start: *t* (5) = 1.645 p = 0.1608; mid *t* (5) = 1.397 p = 0.2212; goal *t* (5) = 9.722 p = 0.002). PVT transients were unrelated to individual approach-avoid decisions if spatial location was ignored (**Figure 2d;** 95% Confidence Intervals). Unlike vSub, PVT activity after choice did not differentiate between approach decisions to stay on the track versus avoid decisions to return to the safety of the start box (**Figure 2e;** 95% Confidence Intervals). Instead, PVT activity selectively increasing during approach–avoid decisions at the goal zone (**Figure 2f;** 95% Confidence Intervals).

How do these vSub and PVT neural dynamics relate to approach–avoid decision-making dynamics? To answer this, we first used HMC to derive LBA decision-making parameters (reward salience, danger salience, decision caution) for each mouse during approach-avoid decisions across the track and then we correlated these with vSub and PVT Ca^2+^ dynamics. We found that vSub Ca^2+^ dynamics were unrelated to the components of approach–avoid decisions (**Figure 2g**) (all R^2^ < 0.20 all p > 0.05). In contrast, there were strong fits between PVT Ca^2+^ dynamics and each of these components (**Figure 2g**) (approach R^2^ = 0.68, *F* (1, 16) = 33.59 p = 0.0001; avoid R^2^ = 0.28, *F* (1, 16) = 5.99 p = 0.0263; caution R^2^ = 0.45, *F* (1, 16) = 13.32 p = 0.0022). Changes in PVT Ca^2+^ dynamics were most strongly associated with the increases in decision caution and reductions in reward salience. Moreover, these decision-making dynamics could be applied within a regression model to accurately predict PVT Ca^2+^ dynamics (**Figure 2h**) (β approach = − 0.69, β avoid = 0.46, R^2^_Adj_ = 0.73, *F* (2, 15) = 23.90 p = 0.0001).

### A prelimbic → paraventricular thalamus pathway controls decision caution in approach–avoidance decisions

These findings show that PVT closely tracks approach–avoidance decisions during motivational conflict but not how it contributes to these decisions. PVT activity co-varied most strongly with the dynamic reductions in reward salience and with increases in decision caution as mice approached the dangerous goal. So, PVT could contribute to these changes in salience (Campus et al., 2019; Zhu et al., 2018), response caution, or both.

To distinguish between these possibilities, we infused an AAV encoding the inhibitory opsin halorhodopsin (eNpHR3.0) (n = 6) or the control enhanced yellow fluorescent protein (eYFP) (n = 6) into prelimbic cortex (PL) (**Figure 3a, S6**). PL is critical for approach-avoid decision-making (Kyriazi et al., 2020; Verharen et al., 2019) and is a major source of excitatory glutamatergic inputs driving PVT neuronal activity (Li and Kirouac, 2012; Otis et al., 2019; Vertes, 2002), so we hypothesized that the role of PVT could depend on this prefrontal input. We implanted optic fibers above PVT allowing us to photoinhibit the PL→PVT pathway (**Figure 3a, Figure S6**). Mice were tested twice under conflict, once in the absence of photoinhibition (Off) and once when the PL→PVT pathway was silenced via 625nm light (On). We photoinhibited the PL→PVT pathway only when mice were at the goal zone, not elsewhere on the track, because fibre photometry had shown that the goal zone was the only location with significant increases in PVT Ca^2+^ transients.

**Figure 3.**
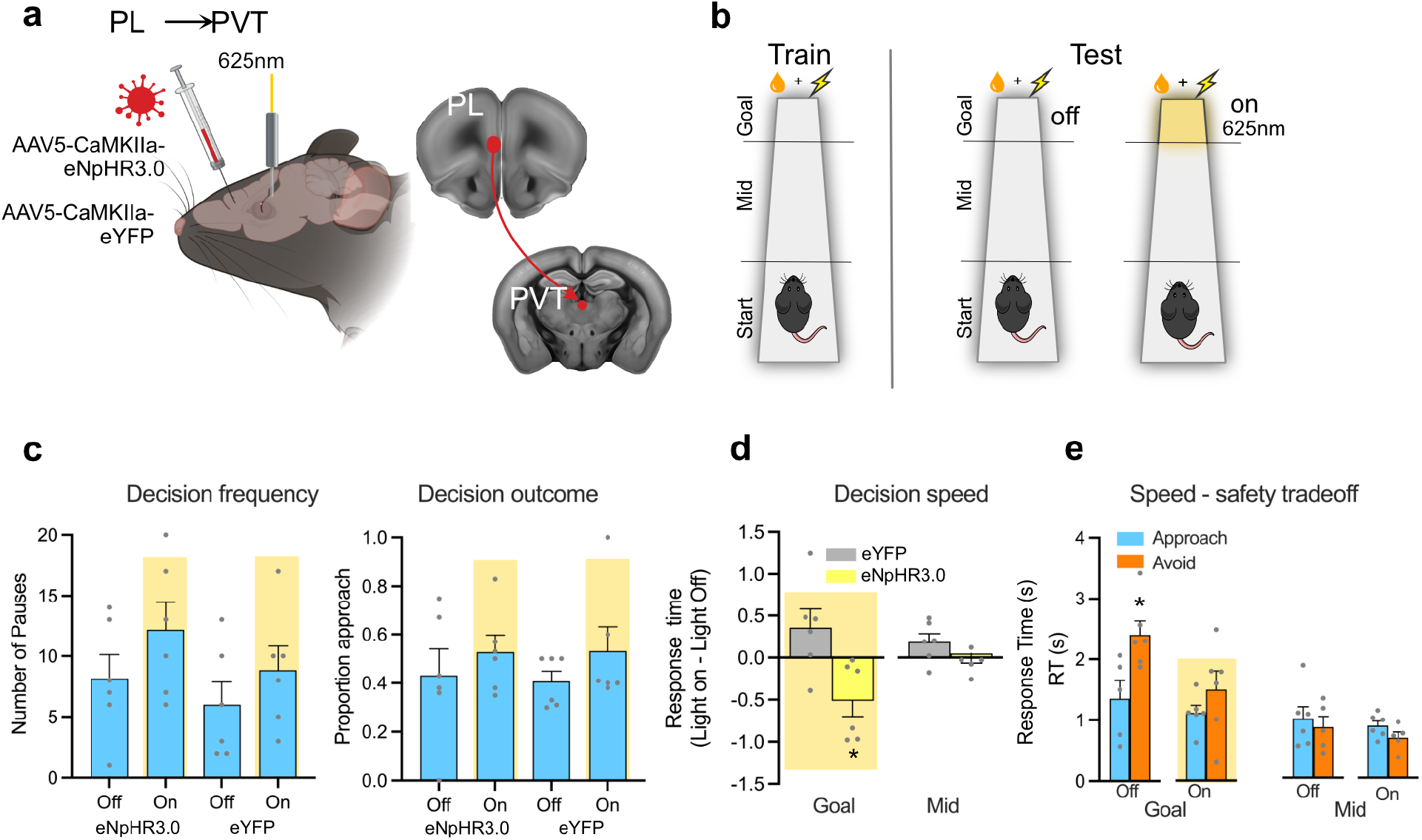
A corticothalamic pathway controls response caution. **a**, AAV encoding the inhibitory halorhodopsin (eNpHR3.0 n = 6) or eYFP (n = 6) was targeted to prelimbic prefrontal cortex and a fiber optic cannula implanted above PVT. **b**, Mice were tested under conflict in the absence (Off) or presence of photoinhibition (On), with photoinhibition delivered only at the Goal zone. **c**, Mean and SEM decision frequency and outcome at Goal zone on tests with (On) and without (Off) photoinhibition. **d**, Mean and SEM RTs at Goal and Mid zones. **e**, Mean and SEM response times at Goal and Mid zones for eNpHR3.0 mice on test without (Off) and with (On) photoinhibition. * p < .05. See Table S3 for detailed results.

If increases in PVT activity are important for dynamic reductions in reward salience, then silencing should prevent these reductions and should bias approach-avoid decisions towards approach decisions at the goal zone. In contrast, if increases in PVT activity are important for dynamic increases in decision caution as danger looms, then silencing should prevent this increase in caution and reduce decision speeds at the goal zone.

Silencing the PL→PVT pathway had no effect on the outcome (Group [eNpHR3.0 v eYFP] x Light [On v Off] interaction: *F* (1, 10) = 0.055 p = 0.819) or frequency (Group [eNpHR3.0 v eYFP] x Light [On v Off] interaction: *F* (1, 10) = 0.135 p = 0.721) of approach-avoid decisions (**Figure 3c; Figure S6**), showing that PVT did not control dynamic alterations in reward or danger salience here (Campus et al., 2019; Zhu et al., 2018). Instead, silencing the PL→PVT pathway hastened decision speeds in the goal zone (**Figure 3d**) (Group [eNpHR3,0 v eYFP] x Light [On v Off] interaction: *F* (1, 10) = 8.862 p = 0.014).

Photoinhibition hastened decision speeds specifically because it abolished the trade-off between decision speed and safety when mice were making approach–avoid decisions in the goal zone (**Figure 3e**) (Light Off: eNpHR3.0 v eYFP: Wilcoxon T = −2.023 p = 0.043; Light On: eNpHR3.0 v eYFP: Wilcoxon T = −0.943 p = 0.345). This was specific to the photoinhibition test (**Figure 3d**), specific to the track location where photoinhibition occurred (Mid zone: no group x light interaction: *F* (1, 10) = 0.046 p = .833) (**Figure 3e**), and there was no effect on the number of visits to the goal zone or on the time that mice spent there (**Figure S6**).

## Discussion

Decisions about pursuing goals in the presence of danger must be made as quickly but as safely as possible. Sequential sampling is a highly efficient, domain-general solution to this problem of fast, accurate decision-making (Forstmann et al., 2016; Shadlen and Shohamy, 2016). We show here that a sequential sampling mechanism explains key empirical features of approach-avoid decisions in mice. This mechanism revealed dynamic changes in salience of reward and punishment and strategic adjustments in decision caution as mice made approach–avoid choices under conflict. So, our findings show that decision-making under fundamental survival conditions in mice shares the same lawful features as speeded decision-making in human (Forstmann et al., 2016) and non-human primates (Shadlen and Shohamy, 2016).

A top-down corticothalamic pathway controls strategic adjustments in decision caution during decision-making under motivational conflict. We show that increases in PVT activity increase the amount of evidence required to reach a decision, thereby increasing the time taken to choose between approaching or avoiding a dangerous goal. This increased evidence threshold drives a trade-off between the speed of choice and the safety of choice. Decisions to avoid danger require more evidence and so take more time. This role for the PVT and the corticothalamic pathway in approach-avoid decision making is distinct from, but complementary to, the well-established roles of corticostriatal circuits in action value and action selection (Hannah and Aron, 2021; Weglage et al., 2021). PVT neurons have extensive, and frequently highly collateralised, projections to these corticostriatal circuits (Dong et al., 2017). So, PVT is well placed to broadcast an evidence threshold across corticostriatal action selection networks to ensure that decision-making under conflict is accurate and safe.

PVT has been implicated in a variety of motivated behaviors but a general account of its function remains elusive (Kirouac, 2015). Many PVT-dependent tasks generate motivational conflict because they involve choices between different, incompatible courses of action (McNally, 2021). These include choices between approach and avoid (Engelke et al., 2021), between different defensive behaviours (e.g., fight versus flight) (Ma et al., 2021), between approaching different sources of reward (e.g., sign versus goal tracking) (Campus et al., 2019), and between persisting with or ceasing behaviour that does not yield reward (Hamlin et al., 2009; Otis et al., 2017). Like the approach–avoidance choices studied here, these choices necessitate trade-off between the speed and outcome of decision-making. So, our finding that PVT controls this trade-off by determining the amount of caution exercised in making a choice provides a mechanism for cognitive control applicable across a range of motivated behaviours and tasks. Moreover, it identifies PVT and its prefrontal cortical input as targets to understand and remediate the deficits in decision caution characteristic of unsafe or impulsive choices.

## Acknowledgments

This work was supported by the Australian Research Council (DP200102576). We acknowledge financial support via the UNSW Major Research Equipment and Infrastructure Initiative and UNSW School of Psychology. We thank Scott Brown for his generous advice and support, as well as Peter Lovibond, Gabrielle Weidemann, Simon Killcross, Fred Westbrook, and Chris Dayas for their comments on this manuscript.

## Author contributions

<u>Conceptualization</u>: G.P.M. <u>Software</u>: M.H. and P.J.R.D.B. <u>Formal Analysis</u>: M.H., P.J.R.D.B., E.Z.M. and G.P.M. <u>Experimentation</u>: E.A.C, E.Z.M., M.H., G.P.M. <u>Writing – original draft</u>: G.P.M. <u>Writing – Review & Editing</u>: All authors. <u>Funding acquisition</u>: G.P.M.

---

**Figure S1.**
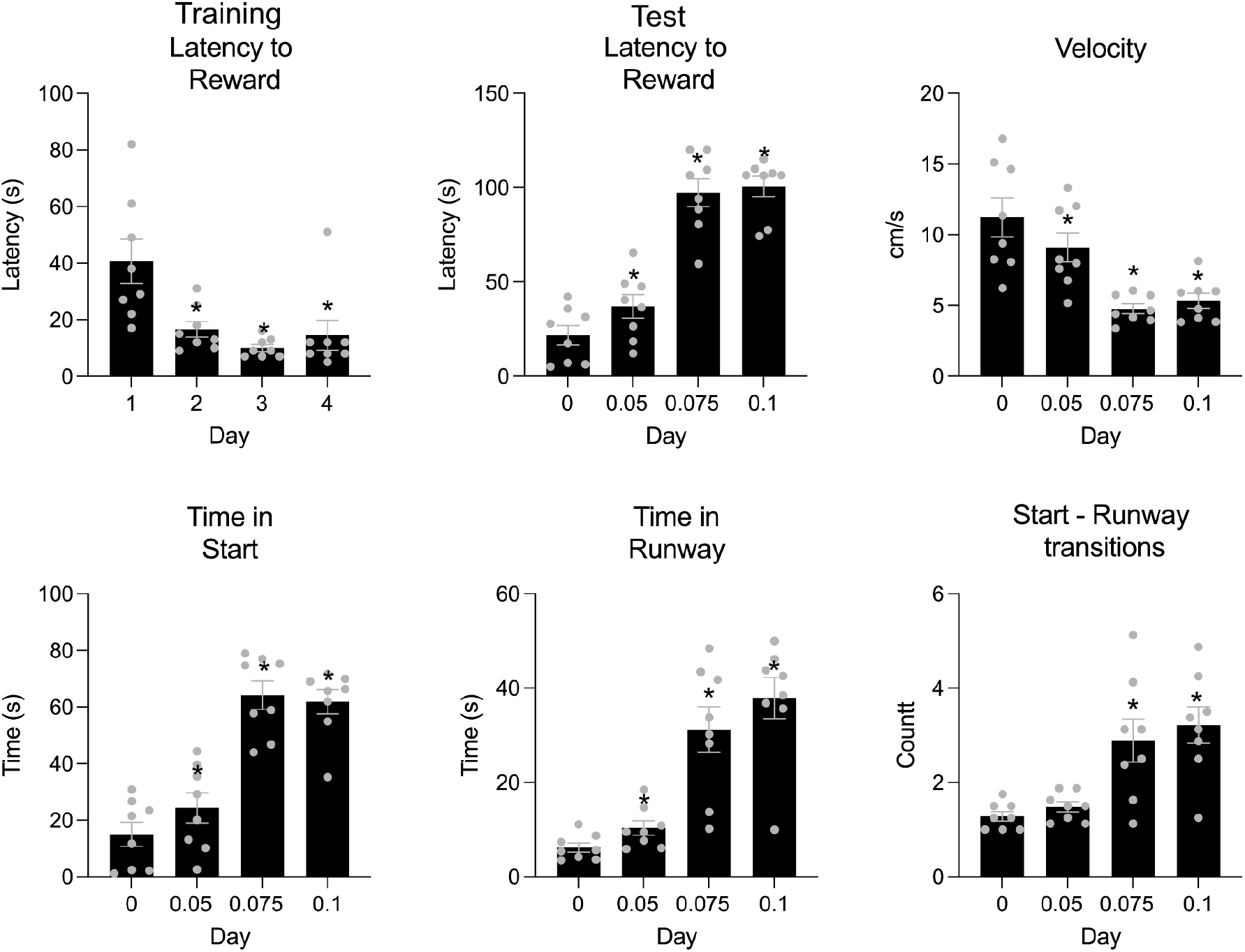
Mean and standard error of the mean (SEM) behavioral measure of conflict. Latencies to obtain reward reduced across training. Conflict increased the time taken to obtain reward, decreased running speeds, increased time in the start box, time on the runway, and the number of transitions between the start box and runway. * p < .05 Table S1.

**Figure S2.**
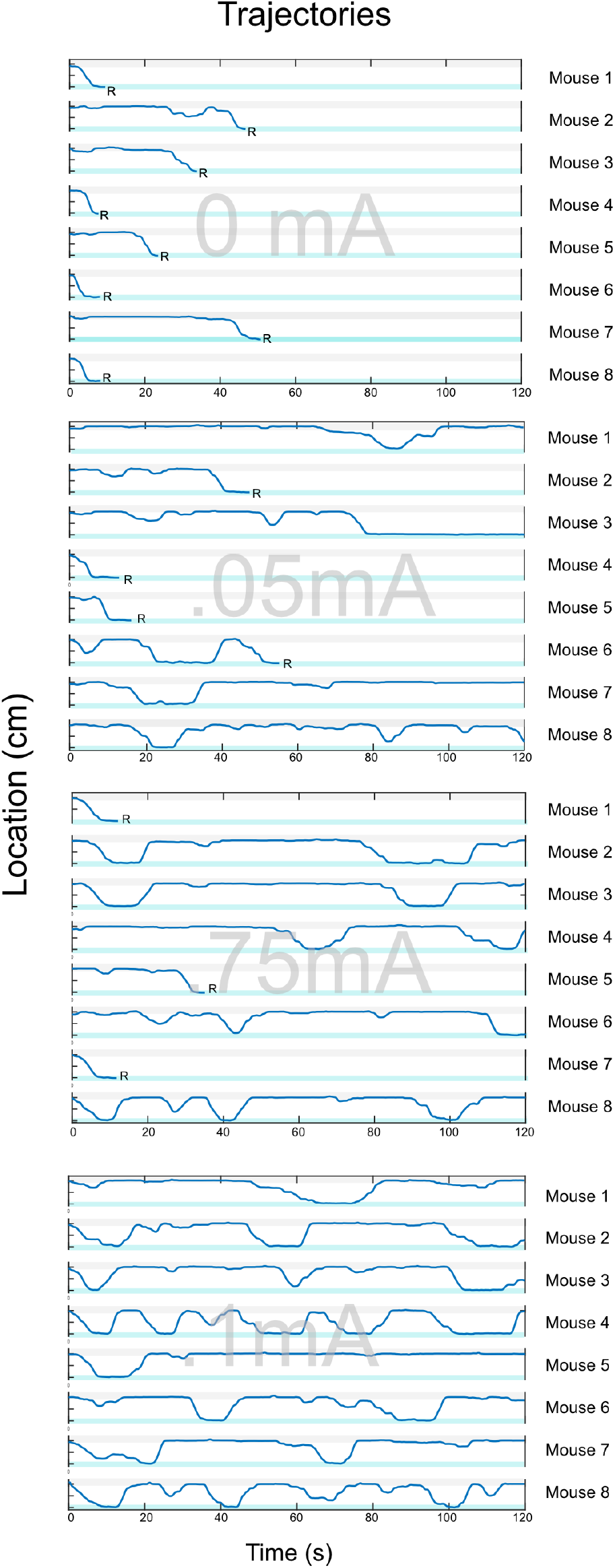
Trajectories of individual mice across conflict at each shock intensity showing emergence of behavioral oscillations. Each row shows an individual animal. Blue shaded regions show start (top) and goal (bottom) boxes. *R* indicates consumption of reward.

**Figure S3.**
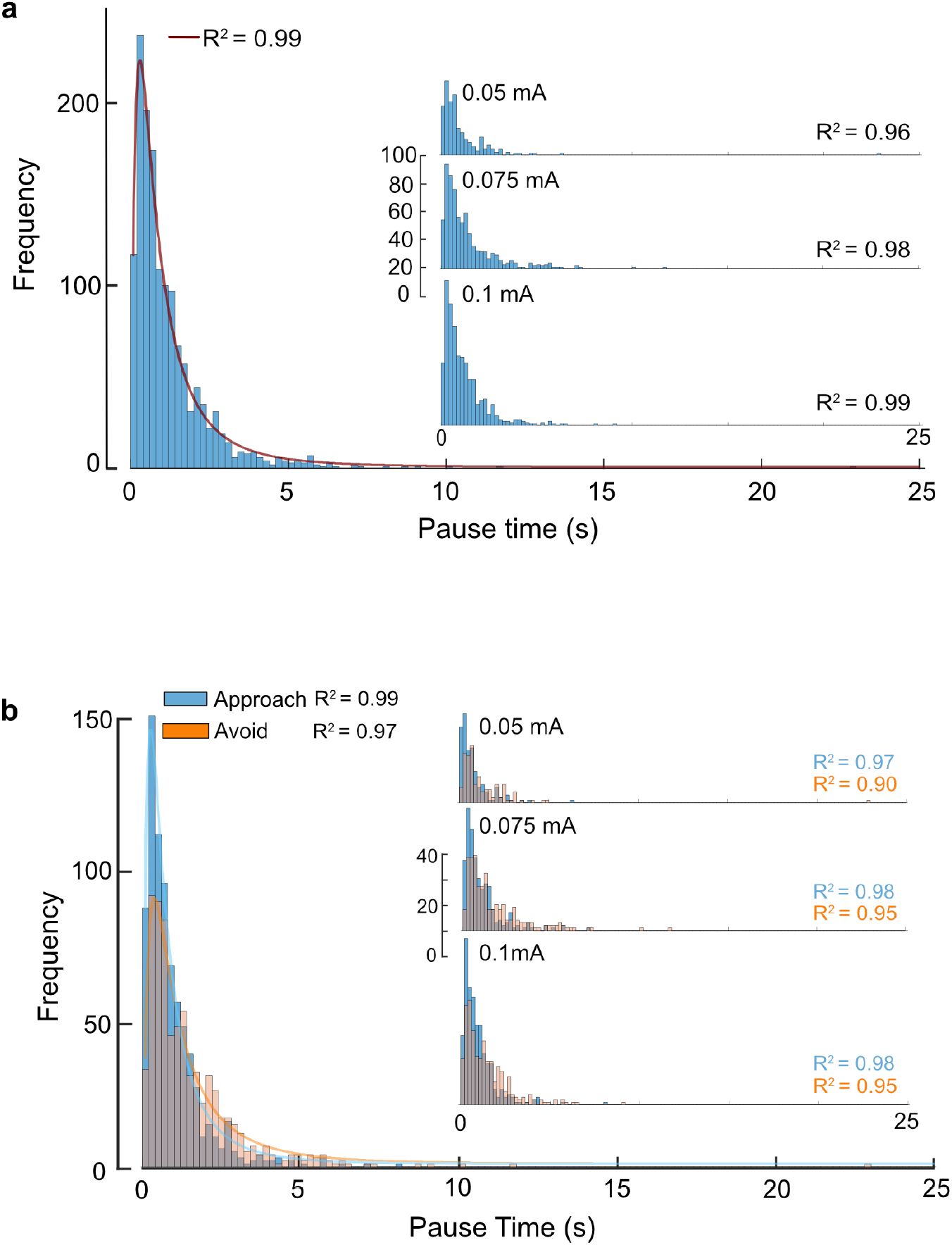
**a**, Frequency distributions and log-normal fits of pause durations by shock intensity. **b**, Frequency distributions and log-normal fits of pause durations by shock intensity and choice outcome. Table S1.

**Figure S4.**
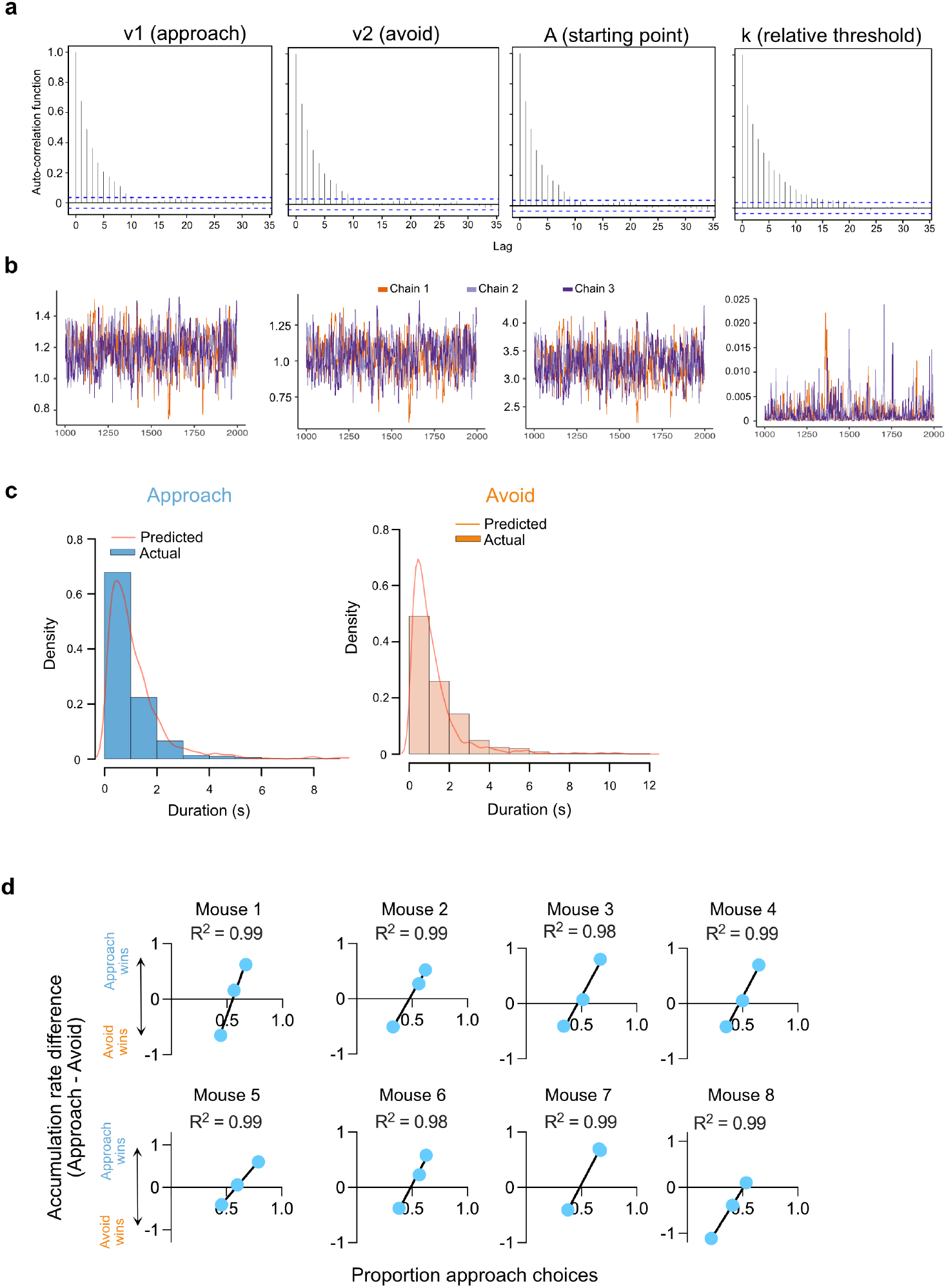
Bayesian parameter estimation of Linear Ballistic Accumulator via Hamiltonian Markov Chain Monte Carlo**. a)** Autocorrelation functions for samples returned by Stan. Autocorrelations dropped to zero at around lags of 10, indicating that the sampler efficiently explored the posteriors for each parameter. **b)** Samples from each chain as a function of iteration showing strong central tendencies, remaining around constant values, with strong overlap between chains, indicating convergence to the posterior distribution. **c)** Posterior predictive check indicating good fit between the model predictions (line) and the observed (histograms) decision times. Warmup=1000; iteration=2000; thinning=1; delta=0.8. Three chains were run to evaluate convergence with a Gelman-Rubin’s criteria of R ̂ < 1.1 and an effective sample size (Neff) > 100. **d)** Predicted choice outcomes (as determined by accumulator with faster rate) and actual choice outcomes across three runway locations for each mouse.

**Figure S5.**
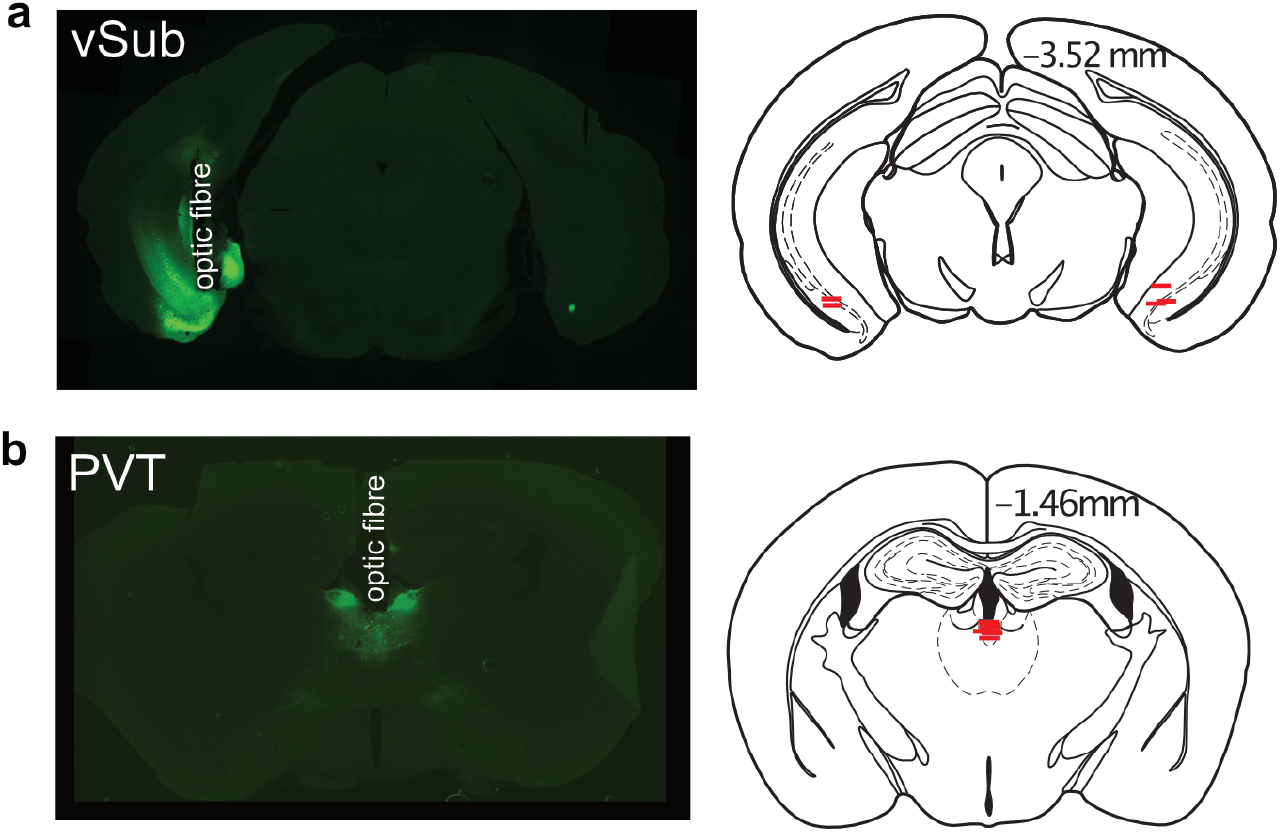
**a**, Representative gCaMP7 expression and fibre optic tip locations in vSub. **b**, Representative gCaMP7 expression and fibre optic tip locations in PVT. Distances in mm from Bregma.

**Figure S6.**
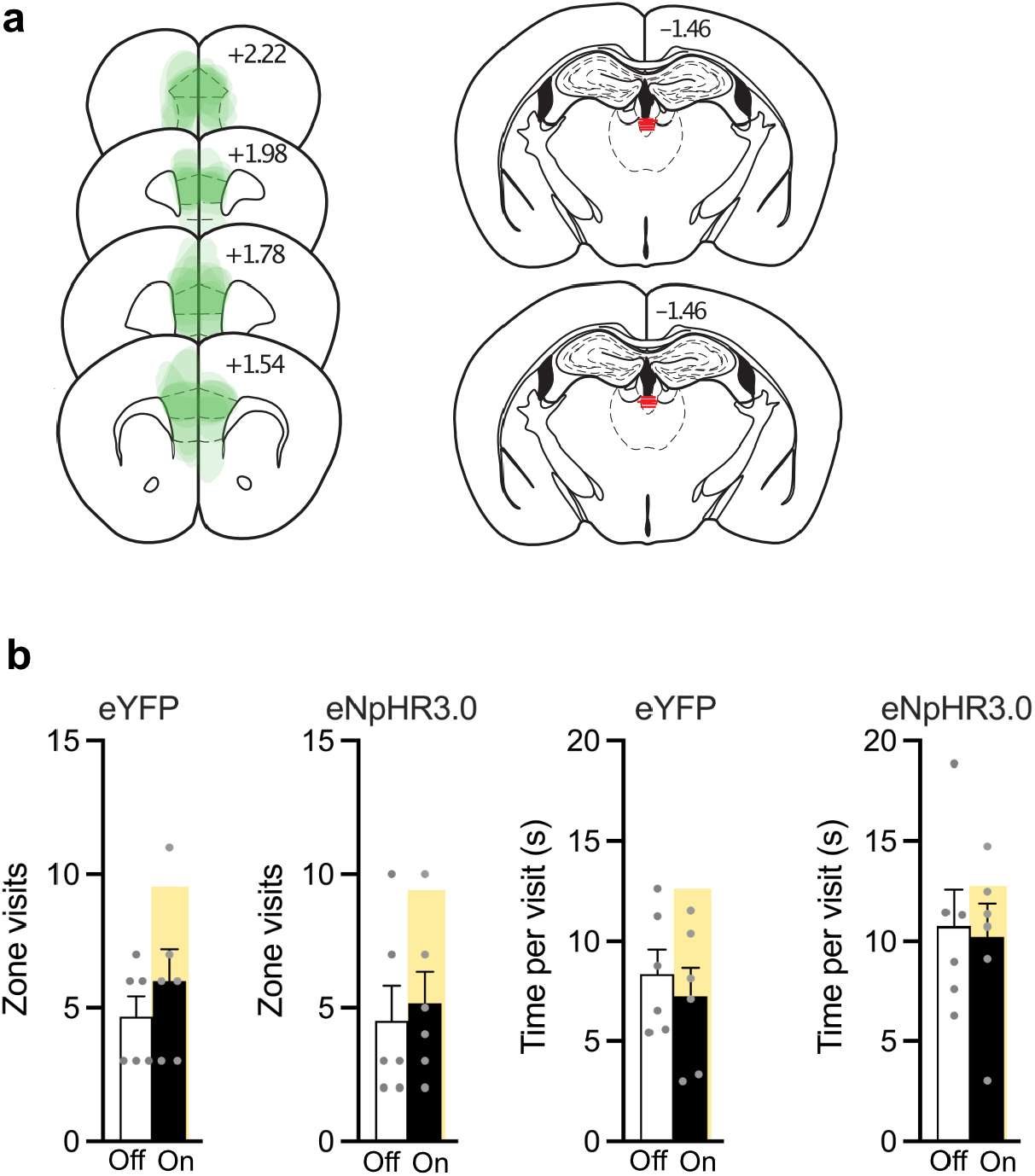
**a**, AAV expression (each mouse is shown at 15% opacity) in prefrontal cortex and fibre optic tip location in PVT. Distances in mm from Bregma. **b**, Mean and SEM number of visits to goal zone on the track and duration of stay per visit. Table S3.

**Table S1.** Statistical Analysis 1 (Fig. 1, Fig S1). a= .05. Significant effects are marked in bold to 4 significant figures.

| Fig. | Test | Dep. Variable | Statistic | p value |
| --- | --- | --- | --- | --- |
| 1c | ANOVA, Dependent means t-test | Pause count | Repeated measures ANOVA Shock intensity<br>$F(3, 21) = 20.38$<br>t-test<br>0 mA: Approach v Avoid $t(7) = 3.230$<br>0.05 mA: Approach v Avoid $t(7) = 2.582$<br>0.075 mA: Approach v Avoid $t(7) = 0.687$<br>0.075 mA: Approach v Avoid $t(7) = 0.070$ | **.0001**<br>**.0144**<br>**.0363**<br>.5137<br>.3201 |
| 1d | Curve fitting | Pause time | Log-normal fit<br>$R^2 = 0.99$<br>$F(1, 122) = 1312$ | **.0001** |
| 1d inset | Regression | Mean and SD pause time | $R^2 = 0.9307$<br>$F(1, 6) = 80.59$ | **.0001** |
| 1e | ANOVA, Dependent means t-test | Pause time | Repeated measures ANOVA<br>$F(2, 14) = 24.86$<br>Dependent means t-test<br>Start v Goal: $t(7) = 4.564$<br>Dependent means t-test<br>t-test: Mid v Goal: $t(7) = 5.555$ | **.0014**<br>**.0026**<br>**.0009** |
| 1f | Dependent means t-test | Pause time | $t(7) = 6.395$ | **.0004** |
| 1h | Dependent means t-test | Reward v Danger accumulation | Mid: $t(7) = 8.134$<br>Goal: $t(7) = 5.555$ | **< .0001**<br>**0.0005** |
| 1i | Dependent means t-test | Caution | Goal v Mid: $t(7) = 4.245$ | **0.0038** |
| S1 | ANOVA, Dependent means t-test | Latency first reward (training) | Repeated measures ANOVA<br>$F(3, 21) = 11.56$<br>Dependent means t<br>0 v 0.05mA $t(7) = 4.193$<br>0 v 0.075mA $t(7) = 8.671$ | **.0020**<br>**.0041**<br>**.0001**<br>**.0001** |
|  |  |  | 0.01mA t (7) = 13.78 |  |
| | | Latency first reward (conflict) | Repeated measures ANOVA<br>$F(3, 21) = 75.21$<br>Dependent means t<br>0 v 0.05mA t (7) = 3.199<br>0 v 0.075mA t (7) = 4.180<br>0.01mA t (7) = 4.923 | **.0001**<br>**.0151**<br>**.0041**<br>**.0017** |
| | | Velocity (conflict) | Repeated measures ANOVA<br>$F(3, 21) = 8.431$<br>Dependent means t<br>0 v 0.05mA t (7) = 4.145<br>0 v 0.075mA t (7) = 6.061<br>0.01mA t (7) = 5.851 | **.0001**<br>**.0043**<br>**.0005**<br>**.0006** |
| | | Time in start box | Repeated measures ANOVA<br>$F(3, 21) = 3.609$<br>Dependent means t<br>0 v 0.05mA t (7) = 3.354<br>0 v 0.075mA t (7) = 10.66<br>0.01mA t (7) = 10.53 | **.0003**<br>**.0122**<br>**.0001**<br>**.0001** |
| | | Time in runway | Repeated measures ANOVA<br>$F(3, 21) = 26.27$<br>Dependent means t<br>0 v 0.05mA t (7) = 3.173<br>0 v 0.075mA t (7) = 4.676<br>0.01mA t (7) = 7.197 | **.0001**<br>**.0156**<br>**.0023**<br>**.0002** |
| | | Transitions | Repeated measures ANOVA<br>$F(3, 21) = 12.73$<br>Dependent means t<br>0 v 0.05mA t (7) = 1.572<br>0 v 0.075mA t (7) = 3.102<br>0.01mA t (7) = 4.508 | **.0027**<br>.1601<br>**.0173**<br>**.0028** |
| S3A | Curve fitting Log Normal | Pause time | Log-normal fit<br>$R^2 = 0.9307$<br>$F(1, 122) = 1312$ | **.0001** |
| S3A | Curve fitting Log Normal | Pause time | 0.05mA $R^2 = 0.96$<br>$F(1, 122) = 186.38$<br>0.075mA $R^2 = 0.98$<br>$F(1, 122) = 187.79$<br>0.mA $R^2 = 0.99$<br>$F(1, 122) = 228.37$ | **.0001**<br>**.0001**<br>**.0001** |
| S3B | Curve fitting Log<br>Normal | Pause time | <u>Overall</u> |  |
| | | | Approach $R^2 = 0.99$ | |
|  |  |  | F (1, 122) = 231.36 | **.0001** |
| | | | Avoid $R^2 = 0.97$ | |
|  |  |  | F (1, 122) = 308.73 | **.0001** |
|  |  |  | <u>0.05mA</u> |  |
| | | | Approach $R^2 = 0.97$ | |
|  |  |  | F (1, 122) = 95.66 | **.0001** |
| | | | Avoid $R^2 = 0.90$ | |
|  |  |  | F (1, 122) = 130.78 | **.0001** |
|  |  |  | <u>0.075mA</u> |  |
| | | | Approach $R^2 = 0.98$ | |
|  |  |  | F (1, 122) = 126.68 | **.0001** |
| | | | Avoid $R^2 = 0.95$ | |
|  |  |  | F (1, 122) = 144.38 | **.0001** |
|  |  |  | <u>0.1mA</u> |  |
| | | | Approach $R^2 = 0.98$ | |
|  |  |  | F (1, 122) = 156.84 | **.0001** |
| | | | Avoid $R^2 = 0.95$ | |
|  |  |  | F (1, 122) = 185.32 | **.0001** |

**Table S2.** Statistical Analysis 2 (Fig. 2). a= .05. Significant effects are marked in bold.

| Fig. | Test | Dep. Variable | Factors | Statistic | p value |
| --- | --- | --- | --- | --- | --- |
| 2b | na | na | na | na | na |
| 2c | ANOVA, Dependent means t-test | df/f<br>Start, Mid,<br>Goal | | <u>PVT</u><br>RM ANOVA $F(2, 15) = 40.62$<br>Start: One-sample vs 0% $t(5) = 1.645$<br>Mid: One-sample vs 0% $t(5) = 1.397$<br>Goal: One-sample vs 0% $t(5) = 9.722$<br><u>vSub</u><br>RM ANOVA $F(2, 12) = 7.51$<br>Start: One-sample vs 0% $t(4) = 3.620$<br>Mid: One-sample vs 0% $t(4) = 9.147$<br>Goal: One-sample vs 0% $t(4) = 6.413$ | **< .0001**<br>.1608<br>.2212<br>**.0002**<br>.0077<br>**.0224**<br>**.0008**<br>**.0030** |
| 2d | Confidence interval | df/f<br>0 – 5 s<br>pause onset | - | 95% Confidence interval | na |
| 2e | Confidence interval | df/f<br>0 – 5 s<br>pause offset | - | 95% Confidence interval | na |
| 2f | Confidence interval | df/f<br>0 – 5 s<br>pause onset | - | 95% Confidence interval | na |
| 2g | Simple regression | df/f<br>0 -5 s pause<br>onset,<br>approach<br>and avoid<br>accumulation<br>rates,<br>caution | <u>vSub</u><br>Approach<br>Avoid<br>Caution<br><u>PVT</u><br>Approach<br>Avoid<br>Caution | <u>vSub</u><br>$R^2 = 0.02$, $F(1, 13) = 0.27$<br>$R^2 = 0.19$, $F(1, 13) = 3.06$<br>$R^2 = 0.03$, $F(1, 13) = 0.37$<br><u>PVT</u><br>$R^2 = 0.68$, $F(1, 16) = 33.59$<br>$R^2 = 0.28$, $F(1, 16) = 5.99$<br>$R^2 = 0.45$, $F(1, 16) = 13.32$ | 0.6108<br>0.1037<br>0.5554<br>**< 0.0001**<br>**0.0263**<br>**0.0022** |
| 2h | Multiple regression – stepwise | df/f<br>0 -5 s pause<br>onset,<br>approach<br>and avoid<br>accumulation<br>rates | <u>vSub</u><br>Approach<br>Avoid<br><u>PVT</u><br>Approach<br>Avoid | <u>Models</u><br>$\beta_{\text{approach}} = 0.06$<br>$\beta_{\text{avoid}} = 1.26$<br>$R^2_{\text{Adj}} = 0.06$, $F(2, 12) = 1.44$<br>$\beta_{\text{approach}} = -0.69$<br>$\beta_{\text{avoid}} = 0.46$<br>$R^2_{\text{Adj}} = 0.73$, $F(2, 15) = 23.90$ | .8603<br>.8325<br>.2747<br>**< .0001**<br>**.0373**<br>**< .0001** |

**Table S3.** Statistical Analysis 3 (Fig. 3). Alpha level was set at p < .05. Significant effects are marked in bold.

| Fig. | Test | Dep. Variable | Factors | Statistic | p value |
| --- | --- | --- | --- | --- | --- |
| 3c | ANOVA | Pause Freq<br>Decision outcome | Group<br>Off, On | <u>Frequency</u><br>Group main effect: $F(1, 10) = 1.323$<br>Light main effect: $F(1, 10) = 4.626$<br>Interaction: $F(1, 10) = .135$<br><u>Outcome</u><br>Group main effect: $F(1, 10) = .010$<br>Light main effect: $F(1, 10) = 2.976$<br>Interaction: $F(1, 10) = .055$ | .277<br>.057<br>.721<br>.922<br>.115<br>.819 |
| 3d | ANOVA | Response time | eNpHR3.0, eYFP, Light on, Light Off | <u>Goal</u><br>Group main effect: $F(1, 10) = 1.661$<br>Light main effect: $F(1, 10) = .300$<br>Interaction: $F(1, 10) = 8.862$<br><u>Mid</u><br>Group main effect: $F(1, 10) = .901$<br>Light main effect: $F(1, 10) = .756$<br>Interaction: $F(1, 10) = .046$ | .226<br>.596<br>**.014**<br>.365<br>.405<br>.833 |
| 3e | Wilcoxon Signed rank | Response time | Off<br>On | <u>Goal</u><br>Approach v Avoid: $T(5) = -2.023$<br><u>Mid</u><br>Approach v Avoid: $T(5) = 0.677$<br><u>Goal</u><br>Approach v Avoid: $T(6) = 0.943$<br><u>Mid</u><br>Approach v Avoid: $T(6) = -0.105$ | **.043**<br>.498<br>.345<br>.917 |

## EXPERIMENTAL MODEL AND SUBJECT DETAILS

Male C57BL/6J (Australian Resources Centre, Perth, Australia) aged 8 – 10 weeks of age were used. Mice were housed in ventilated racks, in groups of 2 - 4, on corn cob bedding in a climate-controlled colony room maintained on 12:12 hour light/dark cycle (0700 lights on). They had free access to food (Gordon’s Mice Chow) and water until two days prior to commencement of behavioral training when they received 30 min of access to water each day for the remainder of the experiment. Experiments were approved by the UNSW Animal Care and Ethics Committee and performed in accordance with the Animal Research Act 1985 (NSW), under both ARRIVE guidelines and the National Health and Medical Research Council Code for the Care and Use of Animals for Scientific Purposes in Australia (2013).

## METHOD DETAILS

### Surgeries and viral injections

Mice were deeply anaesthetized with 5% isoflurane in oxygen-enriched air after subcutaneous injection of 5 mg/kg carprofen (Rimadyl, Zoetis) and then fixed into a stereotaxic alignment instrument (Model 1900, Kopf Instruments). During surgery, mice were maintained on 1–2.5% isoflurane. Before the scalp incision, a local injection of 0.1 ml Marcaine (0.5%) was made subcutaneously at the incision site. Ophthalmic gel (Viscotears, Alcon) was applied to avoid eye drying. Mice received injections of antibiotic (Duplocillin, 0.15ml/kg of body weight subcutaneously) immediately after surgery.

AAVs (0.5 μl) were delivered using a 30-gauge conical tipped microinfusion syringe (SGE Analytical Science) connected to a UMP3 with SYS4 Micro-controller microinjection system (World Precision Instruments). Ventral subiculum coordinates were: 3.15 mm posterior, 2.75 mm lateral, and 4.7 mm below Bregma. Paraventricular thalamus coordinates were: 1.4 mm posterior, 0 mm lateral and 3.15 – 3.35 mm below bregma. Prefrontal cortex coordinates were 1.94 mm anterior, 0.35 lateral (10°) and 2.2 – 2.5 mm below bregma.

### Behavioural Procedures

Mice were placed into a linear track (120 cm (l) x 15 cm (w) x 40 cm (h) constructed of Perspex. A rail ran above the track to which all fibre optic patch cables were attached. There were three segments. The first 22 cm was a start box, constructed from grey and white Perspex walls and a Perspex floor. The Start box was separated from the remainder of the runway by a removable Perspex door. The next 88 cm was the runway proper structed from white Perspex walls and a white Perspex floor. The Goal box was the last 10 cm and was constructed from grey Perspex walls and a stainless-steel grid floor. The goal box contained a stainless-steel extending 3 cm from the end wall for delivery of liquid reward.

For training, there were 4 trials a day for 4 days. Each trial commenced with removal of the door between the Start box and the runway. 10 ul of 8% sucrose was available from the receptacle in the goal box. The trial ended after 2 minutes or after mice had consumed the sucrose. At the end of the trial mice were returned to the Start box for 30 s until the start of the next trial.

For conflict, the same procedures were used but the grid floor in the goal box was electrified using a 0.05, 0.075, or 0.1mA current. Mice received 1 day of training at 0.05mA, 2 days at 0.075mA, and 2 days 0.1mA.

Mice were tracked (30 fps) via webcam (Logitech C920, 1080p) connected to a computer running EthoVision XT 10 (Noldus Information Technology, Wageningen, Netherlands). The runway was divided into three zones – start box, runway, goal box. Ethovision tracked the x- and y-coordinates of the animal’s centre on the runway. From these coordinates the following variables were computed: time spent in these zones, distance from the goal, and velocity of the centre-point of the animal. These data were imported into MATLAB R2018b (MathWorks) for further analysis.

Before locomotor behavior was analysed, frames with no or poor tracking were identified and replaced using linear interpolation. For each frame, a custom script used the x-centre coordinates to determine whether the mouse was moving (≥ 1.5cm uninterrupted) towards the start box, the goal box, or paused (defined as no movement for ≥ 90ms followed by movement of ≥ 1.5cm).

Only pauses that occurred in the runway (i.e. outside the start or goal box) were considered for analysis and modelling. K-means clustering was used to identify distinct patterns of pausing. Pauses within 5 cm bins were collated and median location, total number of pauses, median pause duration, % approach vs avoid decisions were used as inputs in K-means clustering. Silhouette values for 3 clusters were positive (mean = .694, minimum = .243), indicating location thresholds on the runway at 5 cm from goal and 15 cm from start, generating three zones: Start (0 – 15cm), Mid (15 – 83 cm), and Goal (83 – 88 cm).

#### LBA Modelling

The LBA is the simplest, complete model of choice with an analytical solution. Evidence accumulation in the LBA model depends on five parameters (*v, s, A, b, t0*,) where *v* is the drift rate for each response option (sampled on each trial from a normal distribution with mean vi and standard deviation si), *s* is the between-trial variation in v, *A* is the starting point of the accumulation process (sampled on each trial from a uniform distribution), *b* is the amount of evidence needed to make a response, and *t0* is the non-decision time (perceptual and motor processing) (Brown and Heathcote, 2008). Inferences about decision-making processes from LBA are similar to other sequential sampling models with the key advantage that the LBA has a complete analytical solution and can scale to any number of response options.

Following Annis et al. (Annis et al., 2017), we set *s* to a constant value (1) and assumed accumulation rate priors for each response were truncated normal distributions (mean = 2, SD = 1), a uniform prior non-decision time (0, 1), maximum starting evidence for *A* was a truncated normal distribution (mean 0.5, SD = 1), and determined a relative threshold, *k*, from a truncated normal distribution (mean 0.5, SD = 1), from which we could derive b as k + A. Response caution was then defined as b - A/2. To estimate LBA parameters we used a Hamiltonian Markov Chain Monte Carlo algorithm (warmup=1000; iteration=2000; thinning=1; delta=0.8) to obtain posterior distributions. Three chains were run to evaluate convergence with a Gelman-Rubin’s criteria of R ̂ < 1.1 (Gelman and Rubin, 1992) and an effective sample size (Neff) > 100 (Gelman et al., 2013). All analyses were performed using R (version 4.0.2) (Team, 2017) via R Studio (version 1.3.959) and the RStan package (Hoffman and Gelman, 2014; Team, 2015).

### Fibre Photometry

After reward training (see above), mice received 1 day of conflict training with 0.05mA footshock. They were tested the following day with 0.05mA footshock. During this test, mice were tethered to a single fiber optic patch cable attached to a rail that ran above the linear track, providing unhindered motion. Recordings were performed using Fiber Photometry Systems from Doric Lenses and Tucker Davis Technologies (RZ5P, Synapse). Excitation lights (465 nm Ca2+-dependent and 405 nm isosbestic control signal) emitted from LEDs (LEDC1-B_FC, LEDC1-405_FC; Doric Lenses), controlled via dual channel programmable LED drivers (LEDD_4, Doric Lenses), were channelled into 0.39 NA, Ø400μm core multimode pre-bleached patch cables via a Doric Dual Fluorescence Mini Cube (FMC2, Doric Lenses). Light intensity at the tip of the patch was maintained at 10-30μW across sessions. Ca2+ and isosbestic fluorescence were measured using femtowatt photoreceivers (Newport, 2151). Synapse software controlled and modulated excitation lights (465nm: 209Hz, 405nm: 331 Hz), and demodulated and low-pass filtered (3Hz) transduced fluorescence signals in real-time via the RZ5P. Synapse/RZ5P also received timestamping TTL signals from Ethovision.

### Optogenetics

After reward training (see above), mice received 1 day of conflict training with 0.05mA footshock. They were tested the following two days with 0.05mA footshock. During both tests, mice were tethered to a single fiber optic patch cable attached to the rail that ran above the linear track, providing unhindered motion, and which connected to 625nm LEDs (Doric Lenses Inc., Quebec, Canada) controlled by Ethovision. During one test (Off) there was no optical stimulation. During a second test (On), continuous 625nm optical stimulation (10 – 12 mw/mm^2^ measured at the tip of an unimplanted fibre) was delivered only when mice were located within 8 cm of the goal.

### Histology and Immunohistochemistry

Mice were anesthetized with i.p. injections of sodium pentobarbital (100mg/kg) and perfused with 0.9% saline solution containing 1% sodium nitrate and heparin (5000 IU/ml), followed by phosphate buffer solution (PB; 0.1M) with 4% paraformaldehyde. Brains were extracted, incubated in 20% sucrose solution for cryoprotection, sliced coronally (40μm) using a cryostat and stored in PB solution with 0.1% sodium azide at 4°C.

Fiber placements and AAV expression were determined using fluorescent immunohistochemistry. Brain tissue was washed in PB, incubated in PBT-X solution (10% horse serum [NHS], 0.5% Triton X-100 in PB) for 2 hours, and then incubated in PBT-X solution (2% NHS, 0.2% TritonX-100 in PB) with primary antibody (1:1000 polyclonal rabbit anti-GFP, ThermoFisher Scientific, #A11122) at room temperature for 24hr. Tissue was washed with PB and incubated overnight in PBT-X (2% NHS, 0.2% TritonX-100 in PB) with secondary antibody (1:1000 AlexaFluor 488-conjugate anti-rabbit, ThermoFisher Scientific, # A27034). Tissue was washed with PB and mounted onto gelatinised slides. Slides were left to dry and then cover-slipped. AAV expression and cannula placements were verified using fluorescent microscopy. Animals were excluded from analyses if fiber tip and AAV expression could not be confirmed as co-localized.

Criteria for inclusion in final analysis was correct AAV and/or fibre placements determined after histology.

## QUANTIFICATION AND STATISTICAL ANALYSES

Data in figures are represented as mean ± SEM unless otherwise stated. Group numbers for each experiment are indicated at three locations: 1) under the subheadings of behavioural procedures above; 2) in the main manuscript text; 3) in figure legends.

The full results of statistical analyses are show in Tables S1 – S3. Inferential statistics were based on within-subjects t-tests or Wilcoxon-signed rank tests, ANOVA, curve-fitting, and multiple regression. All analyses were conducted using GraphPad Prism version 8.4.2, Matlab, SPSS 25, and the Psy statistical package.

For fiber photometry, Ca^2+^-dependent (465nm-related) and isosbestic (405nm-related) signals and event timestamps were extracted into MATLAB, and signals during logged disconnections were discarded. The isosbestic signal was linearly regressed onto the Ca^2+^-dependent signal to create a fitted isosbestic signal, and a normalized fluorescence change score (dF/F) was calculated using the standard formula: (Ca^2+^-dependent signal – fitted isosbestic) / fitted isosbestic.

Phasic neural activity change around pauses was determined by collating dF/F around pause events (−5s to +5 sec around pause onset or offset, baselined to −5 to −2.5sec pre-evert). To determine significance of activity change, 95% confidence intervals (CI) around activity kernels were derived via bootstrapping (Jean-Richard-dit-Bressel et al., 2020). Bootstrapped means were obtained by randomly resampling from subject mean waveforms with replacement (1000 iterations). CI limits were derived from 2.5 and 97.5 percentiles of bootstrap distribution, expanded by a factor of √(n/(n-1)). A significant transient was identified as a period that CI limits did not contain 0 (baseline) for at least 1/3secs (low-pass filter window). To assess the relationship between neural activity and LBA parameters, mean dF/F during pauses at the 3 runway locations were calculated per subject and correlated against estimates of LBA parameters per subject for these locations.

